# A dynamic Hedgehog gradient orients tracheal cartilage rings

**DOI:** 10.1101/2023.09.25.559425

**Authors:** Evan P. Kingsley, Darcy Mishkind, Tom W. Hiscock, Clifford J. Tabin

## Abstract

The patterning of periodic stripes during embryonic development generates similar structures that repeat at regular spatial intervals within a tissue. These patterns are often attributed to a Turing-like mechanism, which self-organizes characteristically spaced stripes, but these patterns are predicted to be disorganized. Conversely, well-oriented, parallel stripes are often observed in nature. We investigate this phenomenon during the formation of the cartilage rings that support the amniote airway. We find evidence that a Turing-like mechanism underpins the formation of the repeating cartilage elements. Additionally, *SHH* is transiently expressed in a thin dorsal domain along the length of the developing trachea, resulting in a dorsoventral gradient of Hedgehog activity that recedes over time. Using mathematical modelling, we predict that the spatiotemporal dynamics of the gradient are required to organize the stripes into parallel rings. Comparing *in silico* predictions with experimental SHH manipulations shows that the Hedgehog gradient is essential for proper tracheal cartilage patterning.

## INTRODUCTION

Tracheal and bronchial cartilage provide the architectural support necessary to maintain an open respiratory airway for breathing. However, rather than being a continuous sheath, this structural cartilage is organized as a series of distinct elements, either complete rings (e.g., reptiles, birds), or C-shaped rings surrounding the lateral and ventral portions of the trachea (e.g., mammals). This architecture is essential to provide the balance of rigidity and elasticity necessary to maintain air pressure while simultaneously preventing collapse of the airway, as well as allowing flexibility of the neck. Not surprisingly, defects in the development of the tracheal rings result in severe congenital malformations (Javia et al. 2016).

However, in spite of its anatomical simplicity and physiological importance, the development of this simple alternating pattern of cartilage rings and non-cartilaginous connective tissue remains poorly understood. Numerous genetic studies, primarily in mice, have revealed a number of transcription factors and signaling pathways required for the formation of tracheal cartilage rings. For example, mice deficient for the transcription factor *Sox9* in their developing airway fail to form cartilage rings (Hines et al. 2013; Turcatel et al. 2013), while extreme defects in the morphogenesis of the cartilage rings are seen in both the absence of, and the overexpression of Fgf10 signaling (Tiozzo et al. 2009; Sala et al. 2011). Other signaling systems implicated in regulating airway cartilage formation include the Bmp, Shh and Wnt pathways (reviewed by (Iber and Mederacke 2022). How these various signaling systems are integrated to generate a periodic pattern of chondrogenesis has been explored both experimentally and through modeling. These studies have suggested that a Turing-like mechanism may be fundamental to the establishment of the chondrogenic and non-chondrogenic domains within the trachea (Sala et al. 2011; Kingsley et al. 2018; Iber and Mederacke 2022). However, experimental validation of any specific Turing mechanism has remained elusive in the context of the trachea (Iber and Mederacke 2022). Moreover, even in the context of this relatively simple repeating pattern, there are critical aspects that cannot be explained by a Turing system alone. For example, the orientation of the cartilage rings, circumferential and orthogonal to the long axis of the airway, cannot be explained in this manner, and hence remains a fundamental problem in understanding airway development.

The problem of orienting the rings of airway cartilage is only one manifestation of the broader biological problem of orienting Turing patterns. Turing-like local activation/lateral inhibition systems are a central paradigm for explaining self-organized biological patterning, and have proven to be quite powerful in generating periodic patterns like dots and stripes (Turing 1952; Kondo and Miura 2010). However, while Turing mechanisms form periodic patterns, the patterns they generate are disorganized and variable (Green and Sharpe 2015; Hiscock and Megason 2015b). Yet most of the repetitive patterns in nature one would like to explain through such a mechanism are, to a greater or lesser degree, oriented. This is exemplified by many striped animal pigmentation patterns, such as stripes seen in rodents (Mallarino et al. 2016), squamates (Kuriyama and Hasegawa 2017; Murakami et al. 2018), birds (Haupaix et al. 2018; Bailleul et al. 2019), and fish (Singh and Nüsslein-Volhard 2015; Patterson and Parichy 2019). Yet, how the uniform stripes of these and other similar patterns are oriented is mostly unknown. Previous studies have suggested that forming simple, consistent patterns can, at least in principle, be achieved by invoking additional factors such as gradients that affect the parameters or production of system components, e.g., a gradient of signaling molecules (Green and Sharpe 2015; Hiscock and Megason 2015b). However, there is no clear example of such a mechanism being used to orient Turing-generated patterns in nature. The stripe-like formation of digit primordia in the limb bud was proposed to involve a combination of Fgf and Hox gradients interacting with a Turing system (Sheth et al. 2012; Raspopovic et al. 2014) but recent work suggests that the stripe-like morphology also requires mechanically-induced signaling centers which promote elongation of each individual digit (Parada et al. 2022).Thus, the possible roles for gradients in orienting and shaping Turing patterns in embryos, especially stripes, remain opaque, and an example of how this might actually work in a natural setting remains elusive.

Here, we investigate the formation of cartilage rings in the chick trachea (Fig. 1A,B). We show that the pattern of these periodic elements appears to be set up through a Turing-like mechanism, oriented by a spatiotemporally dynamic gradient of Hedgehog (Hh) signaling. By testing predictions from computational simulations of this system against experimental manipulations, we show that this gradient is essential for proper tracheal cartilage formation.

**Figure 1.**
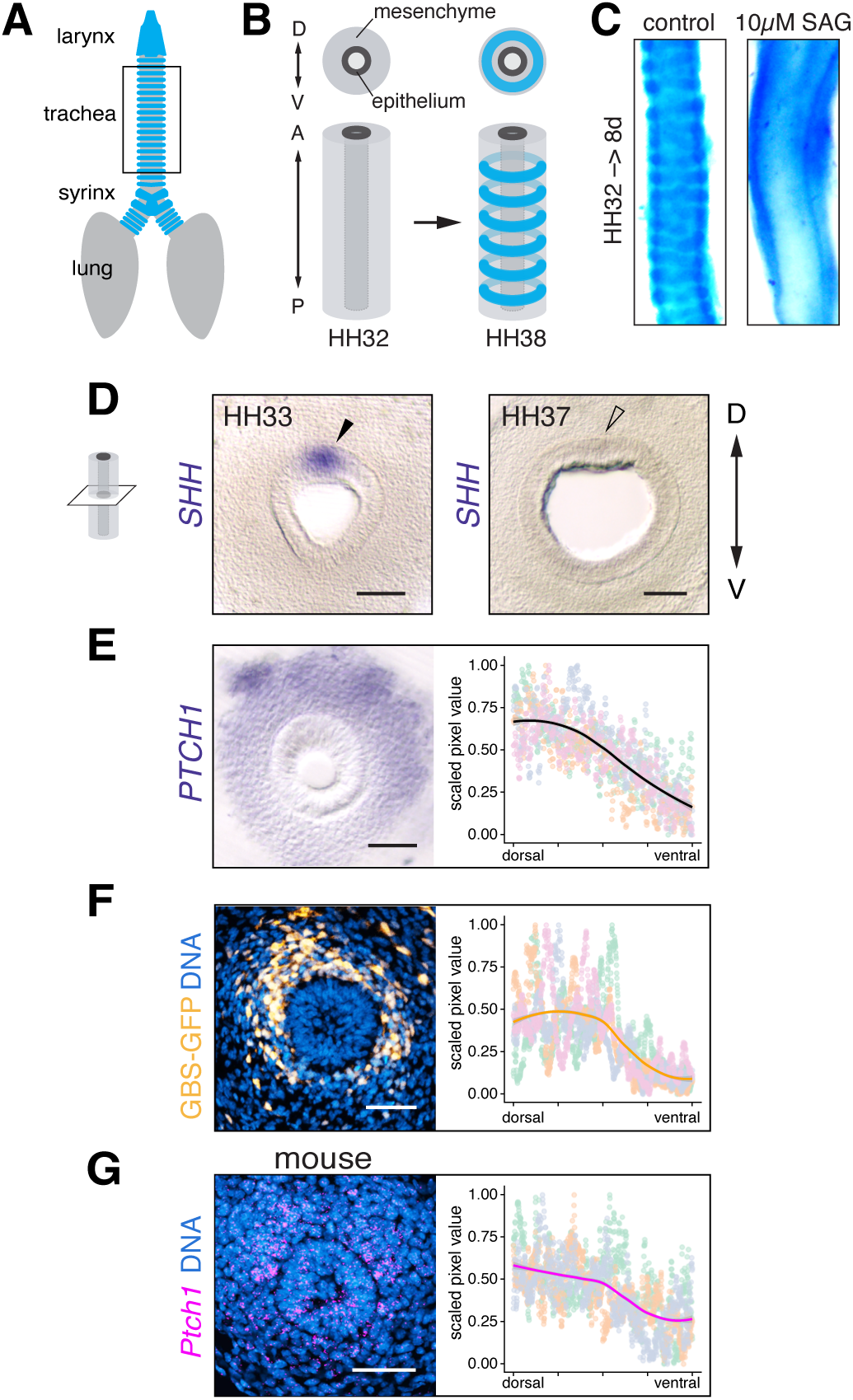
The embryonic chick trachea has a dorsoventral gradient of hedgehog activity. **A.** Schematic of the chick airway, ventral view, showing cartilage in blue. **B.** Schematic showing how the tissue of the trachea (the boxed area in A) begins as an unpatterned tube and goes on to form periodically patterned circular rings of cartilage from the mesenchyme over approximately 5 days of development, from HH32–HH38. **C.** Tracheas explanted at HH32 and cultured for 8 days in the presence of a Hh agonist (SAG) show diffuse alcian blue staining. **D.** ISH on transverse sections chick trachea show *SHH* mRNA expressed in the dorsal epithelium at HH33 (arrowhead, left) and disappears by HH37 (empty arrowhead, right). **E.** ISH shows dorsoventrally graded expression of *PTCH1*, a SHH response gene (center). Plot shows scaled pixel values of *PTCH1* ISH measured on a semicircular path from dorsal-to ventral-most mesenchyme (n = 4 embryos). **F.** RCAN-GBS-GFP infection shows dorsoventrally graded Hh reporter activity in HH33 embryos (n = 4 embryos). Plot is similar to that in E. **G.** *Ptch1* FISH on a transverse section of a mouse E11.5 airway shows a similar dorsoventral gradient. Plot shows scaled pixel values of the *Ptch1* signal, measured as in D, decreasing from dorsal to ventral (n = 3 embryos). For all plots, lines show a loess fit and different point colors represent each embryo. Scale bars = 50µm.

## RESULTS

In a study of chondrogenesis in the chick airway, we uncovered an inhibitory role for Hedgehog (Hh) signaling in tracheal chondrogenesis. Sonic hedgehog (*SHH*) is initially expressed throughout the anteroposterior extent of the developing airway epithelium, but its expression ceases prior to airway cartilage formation. While cultured control airway explants produced cartilage rings similar to those formed *in vivo*, those cultured in the presence of smoothened agonist (SAG), a small-molecule Hh pathway activator, do not form cartilage (Fig. 1C).

To examine the role of SHH activity in the developing chick airway more closely, we examined the *SHH* expression pattern in transverse sections by *in situ* hybridization (ISH). Strikingly, we find that *SHH* expression is actually restricted to the dorsal-most cells of the tracheal epithelium (Fig. 1D). This expression is subsequently lost (Fig. 1D). This localized expression is nonetheless sufficient to promote SHH activity through the dorsoventral extent of the tracheal mesenchyme, as shown by the expression of the SHH target gene *PTCH1* (Marigo et al. 1996) (Fig. 1E). That the observed *PTCH1* expression reflects a domain of active SHH signaling was confirmed by a GLI-binding-site (GBS) Hh reporter (Stamataki et al. 2005; Li et al. 2018) electroporated into the airway primordium (Fig. 1F). Notably, the response to SHH is graded, with the highest level of Hh response (i.e., the highest level of *PTCH1* expression) subjacent to the dorsal *SHH* expression domain.

At an intuitive level, an implication of the Hh activity gradient is that, as SHH production ends and the mesenchymal signal degrades, cells in the ventral mesenchyme should be the first to fall below the threshold level of activity necessary to prevent chondrogenesis, as that is where the signal was already minimal.

Subsequently, the block on chondrogenesis should be lost in a ventral-to-dorsal progression as Hh signaling continues to degrade. This model makes several predictions: First, that tracheal ring chondrogenesis should begin in a small domain in the ventral-most mesenchyme of the trachea, where Hh activity is lowest. When we examine tracheal ring formation by evaluating expression of the early cartilage marker SOX9 and the chondrogenesis marker COL2A1, we indeed find that the chondrogenesis is initiated in the ventral-most aspect of each forming ring (Fig. 2A).

**Figure 2.**
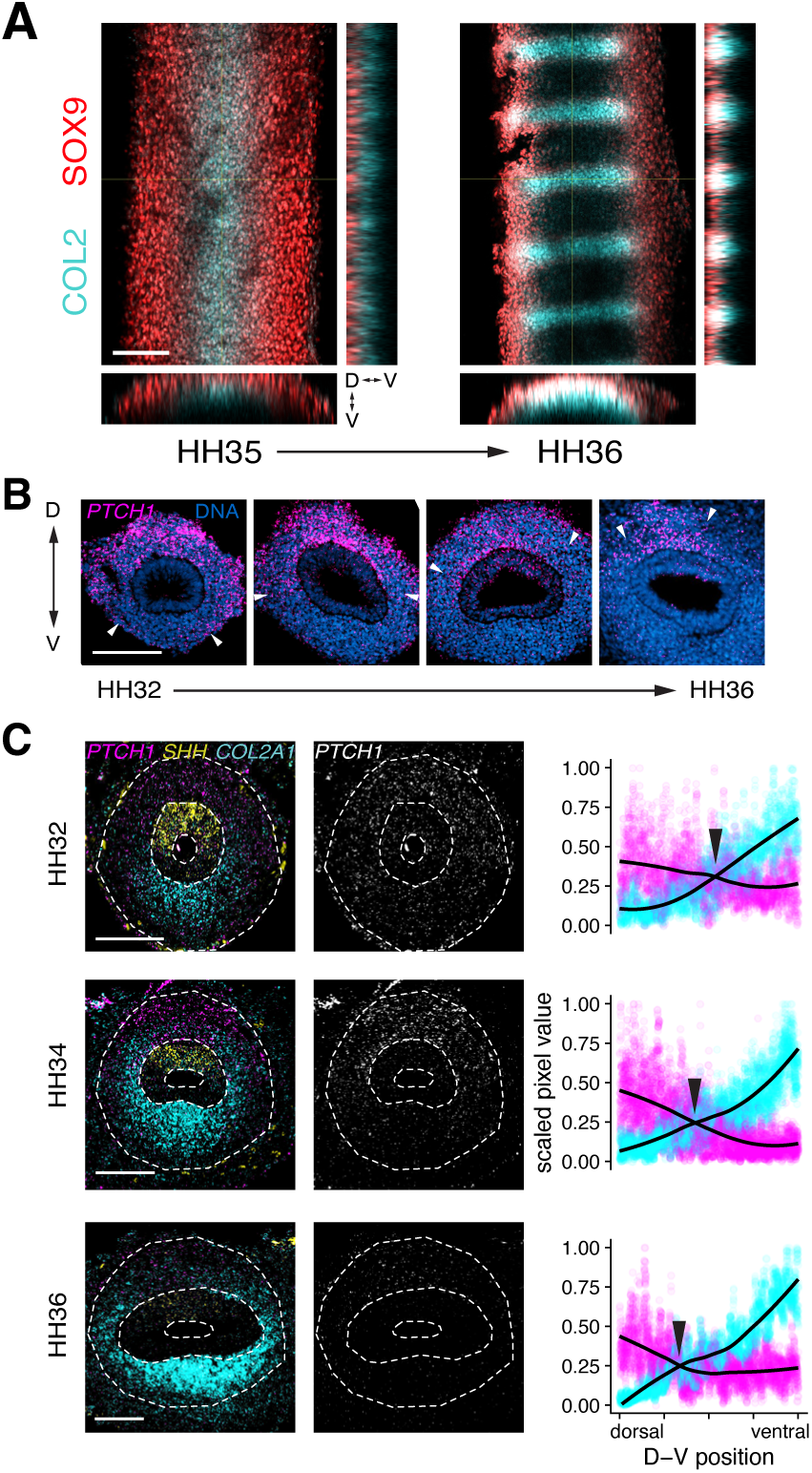
Tracheal cartilage initiates at the ventral surface and expands dorsally as the Hh gradient retracts. **A.** Chick tracheas immunolabeled for cartilage markers show that cartilage elements begin forming at the ventral-most surface of the trachea and expand dorsally. Yellow lines show from where ortho slices are represented on the bottom and right of each image. **B.** Fluorescent *in situ* hybridization shows that the domain of *PTCH1* mRNA expression retracts over time. Arrowheads indicate the ventral edge of the *PTCH1* expression domain. **C.** HCR *in situ* shows that the retraction of the *PTCH1* expression domain through time is concomitant with the expansion of *COL2A1* expression. Plots show scaled pixel values of *PTCH1* (magenta) and *COL2A1* (cyan) expression (n = 3 for each time point). Black lines are loess fits. Scale bars = 100µm.

Second, a careful time series of target gene expression should show that the Hh activity gradient retracts over time. This is indeed the case, as indicated by examining the dynamic changes in *PTCH1* expression as the rings form (Fig. 2B). And third, chondrogenesis should proceed, in accordance with the regression of the Hh gradient, from ventral to dorsal. Simultaneous examination of the temporal dynamics of *SHH*, *PTCH1,* and *COL2A1* expression verified this prediction as well (Fig. 2C). Taken together, these data suggest that the retraction of the dorsoventral SHH activity gradient imposes a spatiotemporal influence on the formation of cartilage rings.

Our data thus suggest how SHH controls the progression of chondrogenesis in a ventral-to-dorsal fashion. The airway cartilage does not form as a sleeve, however, but as a series of discrete parallel elements. How this periodic pattern arises is unknown, but observations are consistent with a Turing-like mechanism. First, the cartilages form essentially simultaneously (Fig. 2A). Additionally, the rings form in a self-organizing fashion, forming even in airway explants in the absence of signals from surrounding tissues (Fig. 1C) (Yoshida et al. 2020). A further indication that the airway rings may be organized through a Turing-like mechanism comes from consideration of the types of irregularities occasionally observed in chicken cartilage rings, and frequently seen in those of the mouse, which resemble the branched and fused patterns generated by Turing-like mechanisms (Fig. 3A) (Vanpeperstraete 1973; Premakumar et al. 2018; Lam et al. 2020). This is especially true of the cartilages observed at the junction between the trachea and bronchi. Simulations show that disordered patterns are often produced where multiple Turing patterns intersect, as at the fork of a Y-shaped geometry (Kingsley et al. 2018). This is precisely what is seen in some, but not all, species at the junction of the more uniform cartilage rings of the tracheal and bronchial regions (Fig. 3B,C) (Heller and Schrötter 1897; Vanpeperstraete 1973). If the tracheal rings are indeed patterned by a Turing-like mechanism, this naturally raises the question of how the Turing-patterned rings achieve their circumferential orientation.

**Figure 3.**
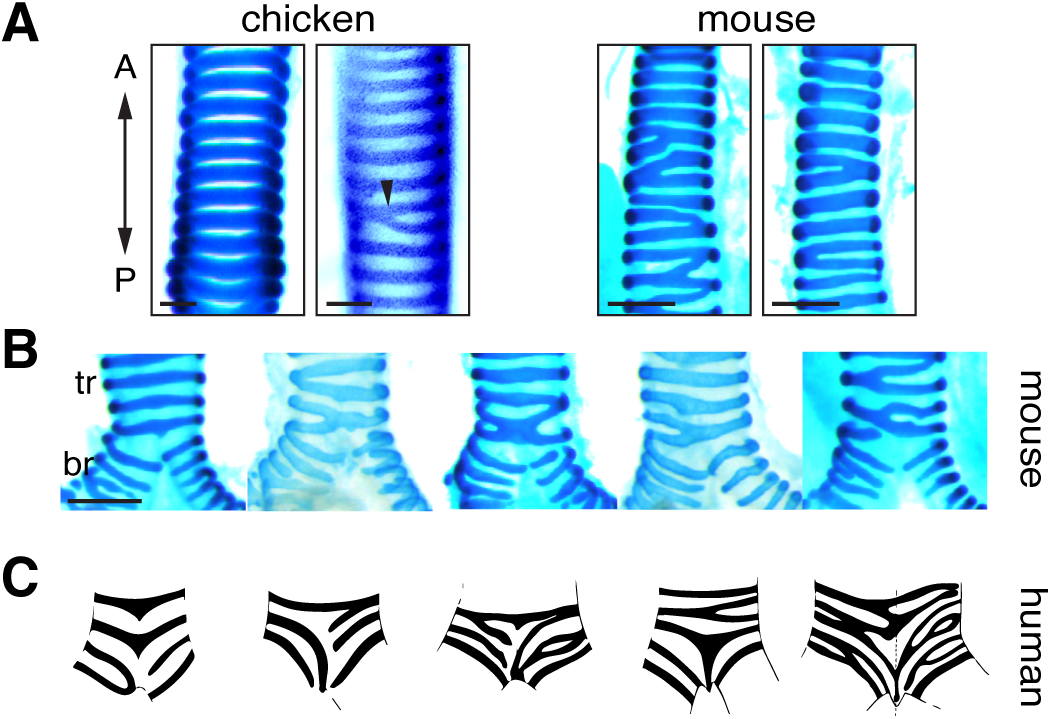
Tracheobronchial cartilage patterns suggest an underlying Turing-like mechanism. **A.** Chick (HH38) and mouse (E18.5) tracheas stained for cartilage showing typical irregularities in the ring pattern for each species. Arrowhead shows an example of a rare branched ring in a chick trachea. **B.** and **C.** Tracheobronchial cartilage of mice (E18.5), stained for cartilage, and human (adapted from illustrations by Heller and Schröter (1879)) showing a variety of cartilage pattern irregularities. Scale bars = 100µm; tr = trachea, br = bronchi.

Prior explorations of pattern orientation have shown that Turing patterns can form parallel stripes in some cylindrical tissue geometries, as on a leopard’s tail (Murray 2003). But this type of tissue geometry only produces circumferential stripes when the tube circumference is on the order of, or smaller than, the pattern wavelength (Fig. S1A). To determine whether the geometry of the trachea could produce an oriented cartilage pattern, we measured airway dimensions at the initiation of chondrogenesis. The ratio of circumference to pattern spacing suggests that geometry alone is insufficient to orient the tracheal cartilage pattern (Fig. S1B), implying that another mechanism is involved.

Previous theoretical work has shown that a gradient affecting parameters of a Turing-like system can generate uniform, parallel stripes that are along the direction of the gradient (Sheth et al. 2012; Hiscock and Megason 2015b). If the initial condensations of the airway cartilage rings are indeed patterned by a Turing-like mechanism, an intriguing possibility is that they could be oriented by the SHH activity gradient, generating circumferential “stripes” of cartilage on a cylinder, i.e., rings. To explore this hypothesis, we formulated a mathematical model of cartilage patterning with two key components: (1) a self-organizing, Turing-like system, which generates regularly spaced stripes of cartilage; and (2) a dynamic Hedgehog gradient, which we hypothesize organizes the otherwise disordered Turing-like system into circumferential cartilage rings (Fig. 4A).

**Figure 4.**
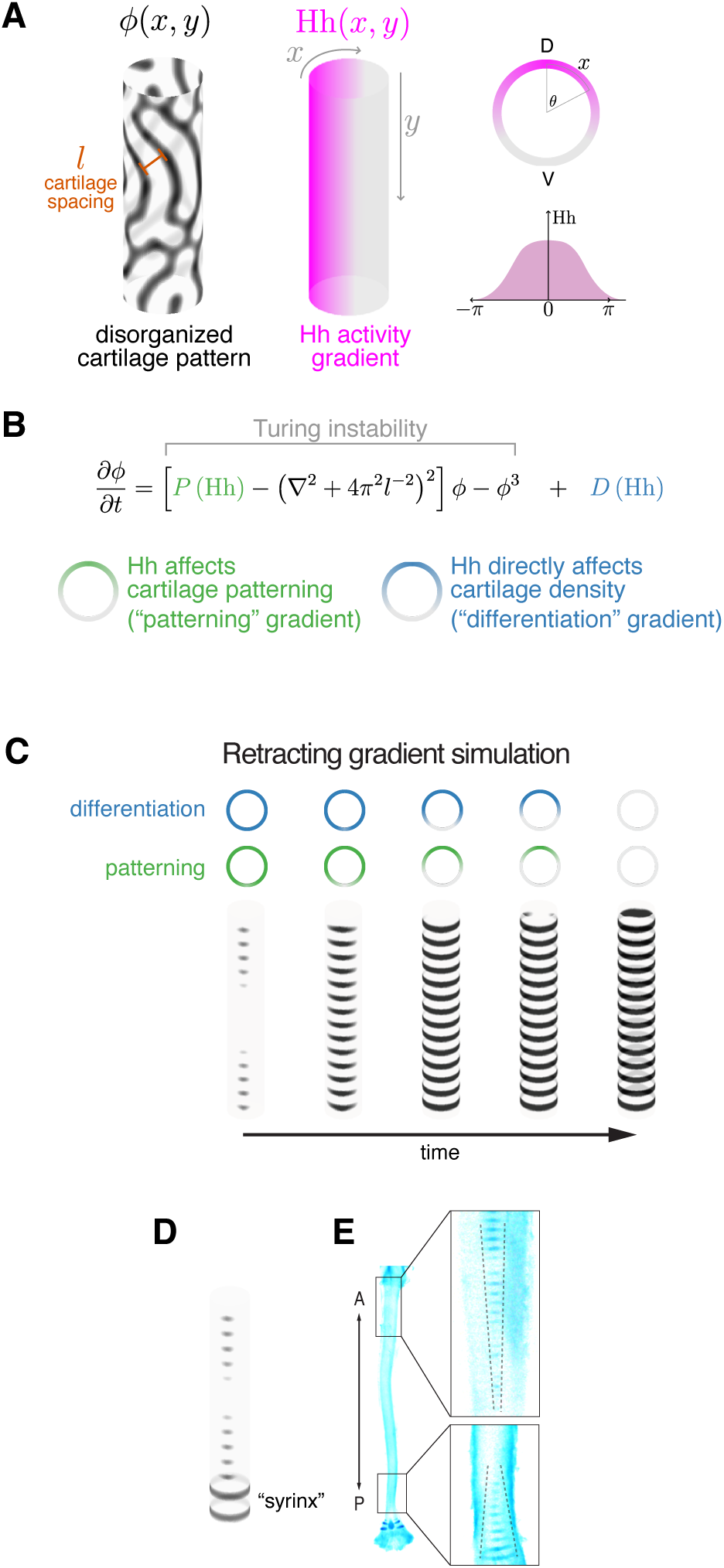
Simulations of tracheal cartilage formation using a generic Turing model coupled to dynamic gradients. **A.** Schematic of a generic Turing model on a cylinder that produces a disorganized pattern (left), onto which we apply a dorsoventral gradient (center and right). **B.** Equation defining the model with “patterning” and “differentiation” terms (see Methods). **C.** Model simulations including retracting, combined patterning and differentiation gradient effects produce a uniform, oriented ring pattern. **D.** Simulations predict a ventral hourglass-shaped domain of initial chondrogenesis due to anterior and posterior boundary effects. **E.** Alcian blue staining shows that cartilage in the chick (HH35) initiates in tapered domains at the anterior and posterior ends of the trachea.

As noted above, whilst there is good evidence that tracheal cartilage patterning is Turing-like, the underlying molecular, cellular and/or mechanical processes remain poorly characterized. Therefore, instead of assuming a specific reaction-diffusion or mechanochemical mechanism, we used a simpler and potentially more generalizable model, the Swift-Hohenberg equation (Fig. 4B), which captures the local activation/long-range inhibition logic common to all Turing-like systems (Cross and Hohenberg 1993). Importantly, previous work has shown that many of the predictions made using the Swift-Hohenberg model – including the effect of gradients – generalize to a diverse range of Turing-like mechanisms, including reaction-diffusion models as well as mechanisms involving cell movement and/or mechanical interactions (Hiscock and Megason 2015a,b). We modified the Swift-Hohenberg equation to explore the effect of a dorsal-ventral gradient in Hedgehog signaling (Fig. 4B). In general, there are two (non-mutually-exclusive) ways that the gradient could impact the Turing dynamics (Hiscock and Megason 2015b). First, Hedgehog could modulate the parameters of the Turing-like patterning system; we refer to this as a *patterning gradient*: P([Hh]). Second, Hedgehog could affect the production/differentiation/density of cartilage directly, independent of its effect on the Turing system; we refer to this as a *differentiation gradient:* D([Hh]). There are a variety of mechanisms that could underpin these terms, depending on the nature of the underlying Turing system. An example of a patterning gradient would be where Hedgehog levels modulate the strength of regulatory interactions between activator and inhibitor molecules in a reaction-diffusion scheme. An example of a differentiation gradient would be where Hedgehog directly repressed the formation of cartilage via inhibition of *SOX9* transcription. In general, we would expect a combination of patterning and differentiation gradients, reflecting the fact that Hedgehog likely affects both P([Hh]) and D([Hh]) terms simultaneously.

Based on our observations of the highly dynamic patterns of Hh activity and its tight coordination with airway chondrogenesis, we allowed the gradients in our model to retract over time (see STAR Methods), hypothesizing that this might be required for reliable pattern orientation. Indeed, simulations revealed that when we added dynamic, receding gradients to our Turing-like model, this resulted in circumferential stripes and reproduced the dorsoventral dynamics of tracheal cartilage formation (Fig. 4C).

Moreover, our simulations recapitulated some unusual edge effects observed at the early stages of tracheal chondrogenesis. We observed transient hourglass-like patterns *in silico* (Fig. 4D), in which cartilage stripes initiated earlier towards the anterior and posterior boundaries, and for a short period were initially longer in these regions, a pattern which is also seen *in vivo* (Fig. 4E). We predict that such patterns are the result of boundary effects which influence the P([Hh]) and D([Hh]) terms at the anterior and posterior extremities (Fig. S2A). In the chick airway, the larynx is located at the anterior end of the trachea and the syrinx sits at the posterior end, raising the possibility that these regions are involved in mediating the boundary effects (Fig. 4E). Consistent with this idea, in species without a syrinx, the lack of a posterior boundary is predicted to produce tapering only in the anterior domain of chondrogenesis; we see precisely this pattern in the alligator airway (Fig. S2B,C).

When we varied gradient parameters in the simulations, we found that ring orientation is robust to variation in the steepness (W) of the gradient, but sensitive to parameters that reflect the strength (P_off_) and range (b_0_) of Hh activity (Fig. 5A) (see STAR Methods for description of gradient parameterization). Intriguingly, variation in these parameters can reproduce many of the types of ring irregularities that are frequently seen in human and mouse airways (compare Fig. 5A with Fig. 3), with different initial conditions giving rise to distinct defect types (Fig. S3). In particular, cartilage elements in the chick trachea rarely branch or fork, while these irregularities are common in mouse and human airways (Fig. 3) (Vanpeperstraete 1973; Premakumar et al. 2018; Lam et al. 2020). We note that the Shh activity gradient appears to be stronger in the chick than in the mouse (Fig. 1E vs. Fig. 1G) raising the possibility that different gradient parameters could cause pattern differences across taxa.

**Figure 5.**
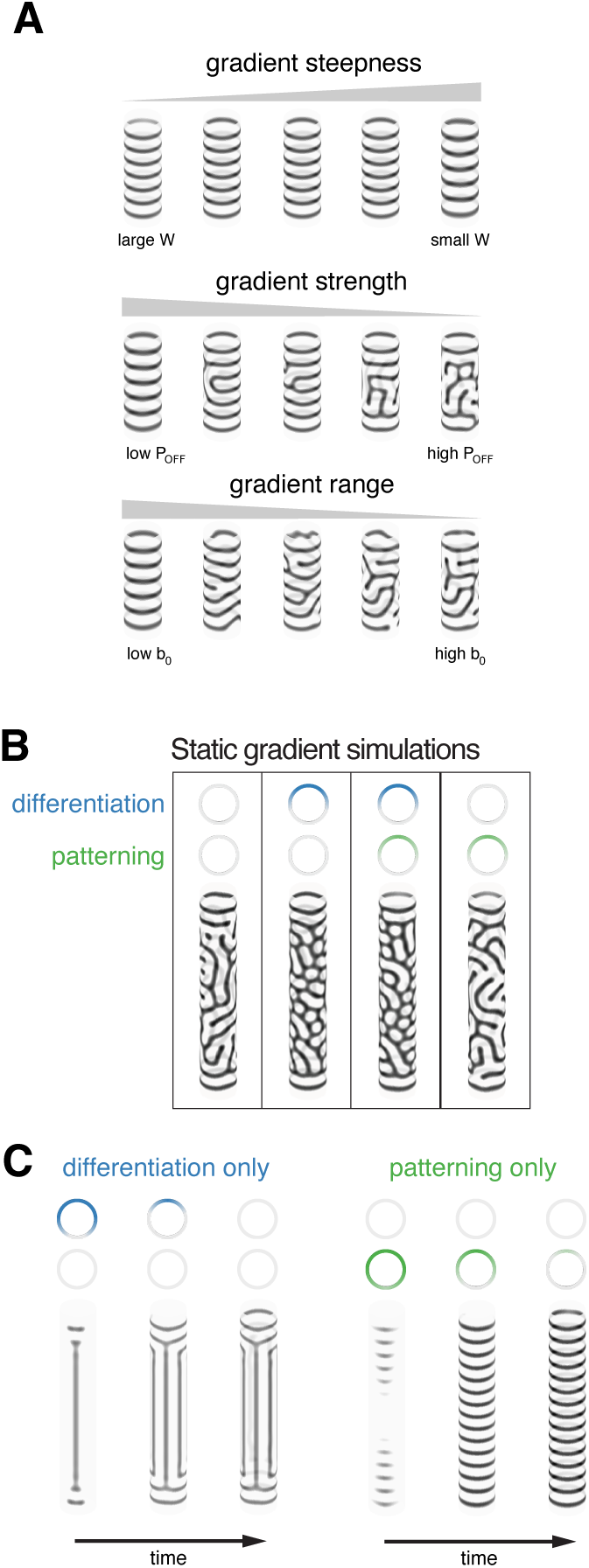
Exploration of varying gradient parameters on pattern orientation. **A.** Simulations of varying model parameters show that ring orientation is robust to changes in gradient steepness (W), but that modifying the strength (P_OFF_) or range (b_0_) of the gradient influences pattern orientation. **B.** Model simulations with no gradient, a single gradient effect, or combined gradient effects predict that static gradients are insufficient to orient ring-like patterns. **C.** Simulation results of models with a single gradient effect show that a retracting differentiation effect alone does not orient rings, while a retracting patterning effect does.

The dynamic nature of the gradient effect is essential: simulations that assume static gradients do not produce oriented rings (Fig. 5B), indicating that the receding dynamics are necessary. Nor does a retracting differentiation gradient, alone, orient rings; a retracting patterning effect is required (Fig. 5C). Theoretical analysis of the Swift-Hohenberg model can explain, rather generally, why a receding patterning gradient is required, a conclusion which is likely to hold for a diverse range of Turing-like systems (see STAR Methods, “Generality of simulation results to other Turing-like systems”). Together, these results are consistent with the idea that dorsoventral retraction of a tracheal Hh activity gradient orients the cartilage ring pattern. This dynamic gradient mechanism is, to our knowledge, a previously undescribed way to reliably orient stripes in a Turing system.

We modeled a generic Turing system because the nature of the hypothesized patterning mechanism is unknown. To show that a spatiotemporally dynamic gradient can orient stripes in a specific type of Turing model in the same way as our generic model, we provide an example of a putative reaction-diffusion model (Fig. S4A). Indeed, the receding gradient can orient rings in this model as in the generic model (Fig. S4B,C). While not aiming to propose any specific molecules, this result shows that our model is applicable to more explicitly defined Turing systems.

To assess the biological plausibility of the retracting gradient model, we sought to perturb the spatiotemporal dynamics of Hh activity both *in silico* and in the chick embryonic airway. Because simulations of the patterning-only model orients rings similarly to the two-effect, differentiation/patterning model, we simulated perturbations of both. Simulations predict that prolonging Hh activity in a restricted region of the airway would block cartilage formation in the vicinity of the increased activity and give rise to mild reorientation defects near the perturbation; this result is robust to variation in the nature of the gradient effect (Fig. 6A). To test this prediction, we explanted chick airways at HH34, when the native SHH activity has partially retracted, and electroporated a plasmid that constitutively expresses chicken *SHH* (pCAGGS-cSHH) into the dorso-lateral epithelium. After 5 days of culture, we saw that mesenchyme adjacent to the electroporated cells fails to produce cartilage condensations (4 of 4 airways) and, although explanted airways produce somewhat irregular cartilage elements as a consequence of the culture conditions, the elements near the perturbation appear to be slightly reoriented compared to the control airways (3 of 4 airways) (Fig. 6B). We then addressed the complementary prediction, in which Hh activity is prematurely reduced. Model simulations predict that systemically blocking Hh activity when the gradient is partially retracted will produce ectopic, disorganized dorsal cartilage condensations that may fuse, depending on the nature of the gradient effect (Fig. 6C). When we explanted airways partway through gradient retraction, at HH34, and treated them with cyclopamine (5µM), we saw that the treated airways produced disorganized, fused dorsal cartilage elements when compared with control airways (3 of 3 airways) (Fig. 6D), which matches the prediction of the two-effect model. This is counter-intuitive: both untreated and cyclopamine-treated tracheas will lose Hh activity, but our results imply that the history of spatiotemporal dynamics is required for proper ring formation.

**Figure 6.**
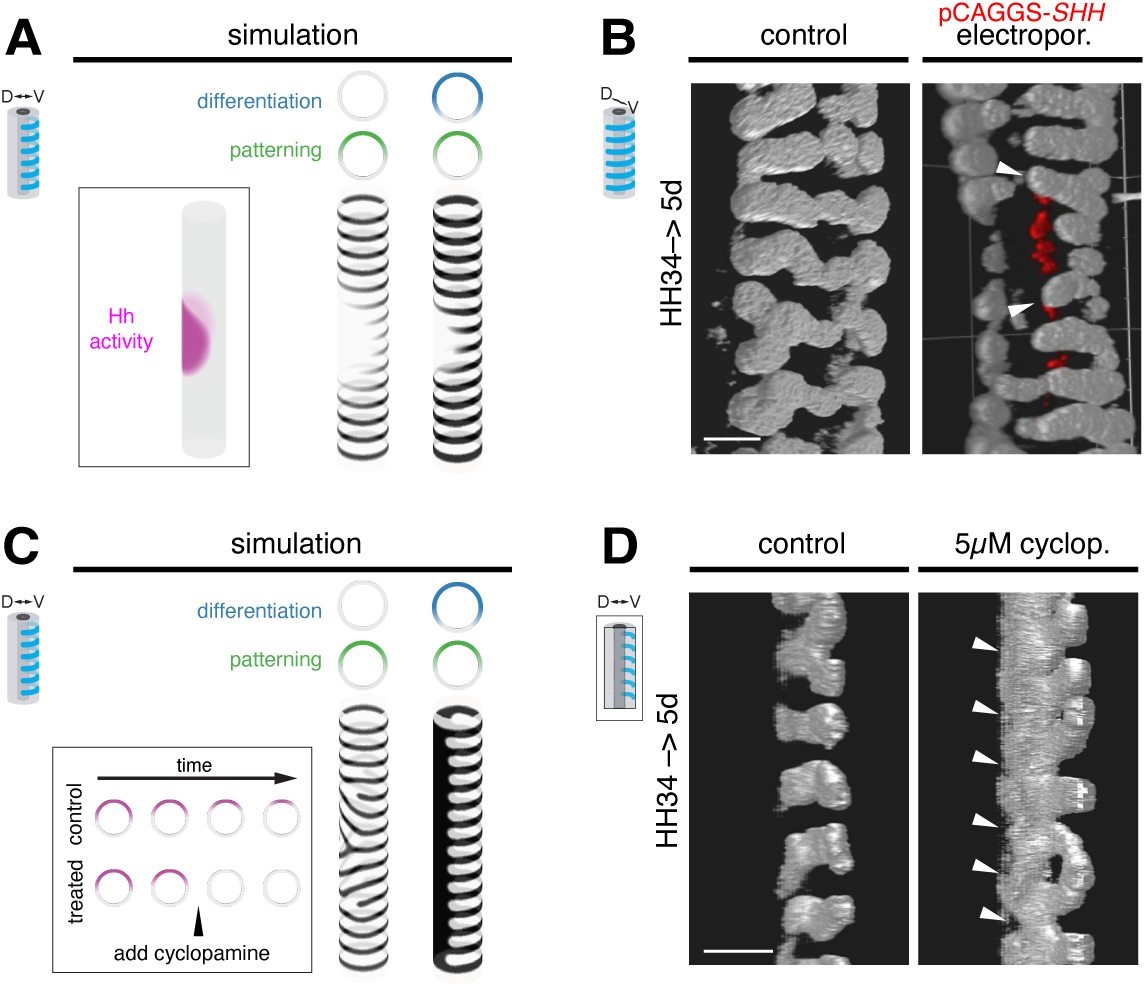
Simulated gradient perturbations predict explant culture outcomes. **A.** Simulations predict that a persistent domain of Hh activity (box) will produce an unpatterned area with a somewhat misoriented pattern at the domain edge. Models with either a retracting patterning-effect gradient (left) or with combined patterning and differentiation effects (right) predict this outcome. **B.** 3D reconstructed cartilage (gray) immunostains of explanted chick airways. Electroporation into the airway epithelium of a plasmid constitutively expressing the chick *SHH* sequence (red) locally inhibits cartilage formation and produces misoriented cartilage elements (arrowheads), resembling the model predictions. **C.** Simulations predict that systemically inhibiting Hh partway through gradient retraction (box) would either partially reorient rings (with a patterning-only gradient effect) or cause cartilage fusions (with both gradient effects). **D.** 3D reconstructed cartilage (gray) immunostains, cutaway view of the left side of airway explants. Cyclopamine-treated airways produce fused cartilage elements, resembling the 2-effect gradient model prediction. Scale bars = 100µm.

We describe SHH having an inhibitory effect on tracheal chondrogenesis. While this is consistent with the way the loss of Hh activity permits differentiation during morphogenesis of other organs (Hebrok et al. 1998; Fagman et al. 2004; Grevellec et al. 2011; Rowton et al. 2022), it stands in contrast to genetic data showing that SHH activity is required for airway cartilage formation in the mouse (Miller et al. 2004; Park et al. 2010; Sala et al. 2011). This seeming contradiction might be explained by a critical difference in the architecture of the tracheal cartilage elements in birds vs. mammals. In birds, the cross section of each tracheal cartilage is a complete ring, whereas in mice and other mammals, the rings are C-shaped, with the most dorsal aspect of the trachea composed of smooth muscle (SM) rather than cartilage. Recent work has shown that SM is required for proper cartilage patterning in the developing mouse airway (Young et al. 2020), which is evidently not true for birds as their tracheas lack smooth muscle (Klingler 2016). Being the dorsal-most tissue, we reasoned that tracheal SM in the mouse may require SHH activity. Indeed, Hh activity is present in a gradient in the mouse trachea (Fig. 1G), and *Shh*-deficient mice fail to properly form the SM of the dorsal trachea (Fig. S5) (3 of 3 +/*Shh-cre* heterozygotes had fully formed dorsal SM; 3 of 3 *Shh-cre*/*Shh-cre* homozygotes did not). Thus, SHH may play multiple roles in forming the supports for the mammalian airway: First, by inducing SM, which is itself essential for robust cartilage patterning, SHH may be indirectly required for proper chondrogenesis, partially explaining the loss-of-function phenotype. And a second, later role for SHH may be to influence the cartilage pattern in a ventrodorsal progression as in the chick. Whether this plausible model is true, and more generally understanding the more complex formation of the airway cartilage in mammals, will require further investigation.

Taken together, our results add further weight to the proposal that the underlying stripe-like chondrogenic pattern in the developing airway is established through a still unspecified Turing mechanism. However, achieving the reliable parallel orientation of the cartilage rings requires an additional input provided by the dynamic dorsoventral gradient of SHH activity.

## DISCUSSION

Despite parallel, periodic stripe patterns being common in animal development, there is much debate about how these patterns form. We have extended prior gradient models of Turing pattern orientation, showing that a spatiotemporally dynamic gradient of SHH activity exists in the embryonic chicken trachea and that the dynamic nature of the gradient is essential for proper airway chondrogenesis. We show that a spatially restricted *SHH* expression domain in airway epithelium produces a dorsoventral gradient of Hh activity in airway mesenchyme. This gradient retracts as *SHH* expression ceases, and this retraction coincides with the dorsal expansion of the cartilage rings. A computational model of an underlying Turing system produces parallel rings if we allow the pattern-generating mechanism to be affected by the dynamic gradient. When we perturb the Hh activity gradient, the airway cartilage pattern is disrupted in ways that are predicted by model simulations. Together, these data provide evidence that a previously undescribed retracting gradient mechanism orients a Turing system to reproducibly form parallel rings of airway cartilage.

There is much about the action of the tracheal SHH gradient that remains unknown. We observe that a model incorporating a dynamic gradient more reliably orients stripes than one with a static gradient. This suggests that, for a context in which producing uniform parallel stripes is optimal, a dynamic gradient may be favored by natural selection to orient rings. Our data imply that changing a static gradient to a receding gradient could be as simple as shutting off production of a secreted ligand that is produced by a subset of cells; is that the only mechanism by which the gradient strength, shape, and retraction speed are regulated? Indeed, prior work in mice has identified a large number of factors that affect the development of tracheal cartilage (Iber and Mederacke 2022), suggesting that other factors could be involved in regulating the gradient and thus the cartilage pattern.

We assume that an underlying Turing system defines cartilage ring periodicity, but what might this system consist of? Interactions among the multiple signaling pathways, including MAPK/ERK, BMP, and WNT/B-catenin, in addition to Hh, that have been shown to influence tracheal cartilage development in mice have not been explored in the context of patterning (reviewed by (Iber and Mederacke 2022)). Recent work in mice suggests that *Col2a1*-expressing cells move and aggregate during tracheal development (Young et al. 2020) and that myosin contractility is essential for tracheal cartilage formation (Yoshida et al. 2020), which may indicate a role for cell-level patterning. Finally, tissue mechanical properties of the epithelial and mesenchymal layers, e.g., differential stiffness or growth, could produce periodic patterns (as in (Shyer et al. 2013; Marin-Riera et al. 2018)). Here, we model the interaction of the dynamic gradient with the parameters of a Turing-like system but knowing the nature of the underlying patterning mechanism will be key to understanding how the gradient influences that system.

How generally applicable could the gradient model of pattern orientation be? A number of mechanisms have been proposed to orient Turing-like systems into stripes, including orienting to an initial condition (Nakamasu et al. 2009), tissue or diffusion anisotropies (Shoji et al. 2002; Das 2017), extreme tissue geometries creating boundary conditions (Murray 1988), and interactions with gradients (Sheth et al. 2012; Hiscock and Megason 2015b). Proposed interactions of Turing systems with signaling gradients, in particular, vary widely and may be common.

## STAR METHODS

### Key Resources Table

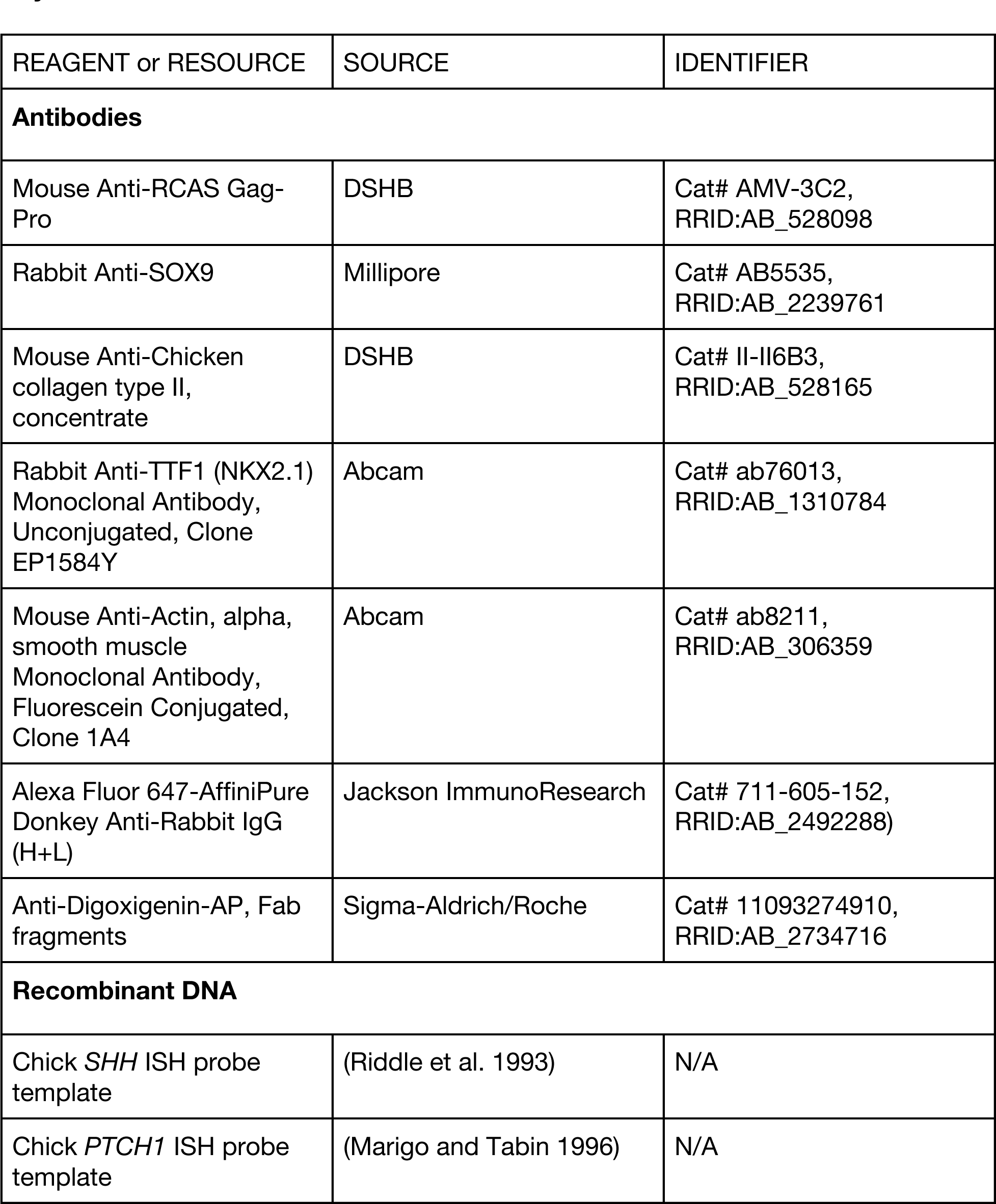
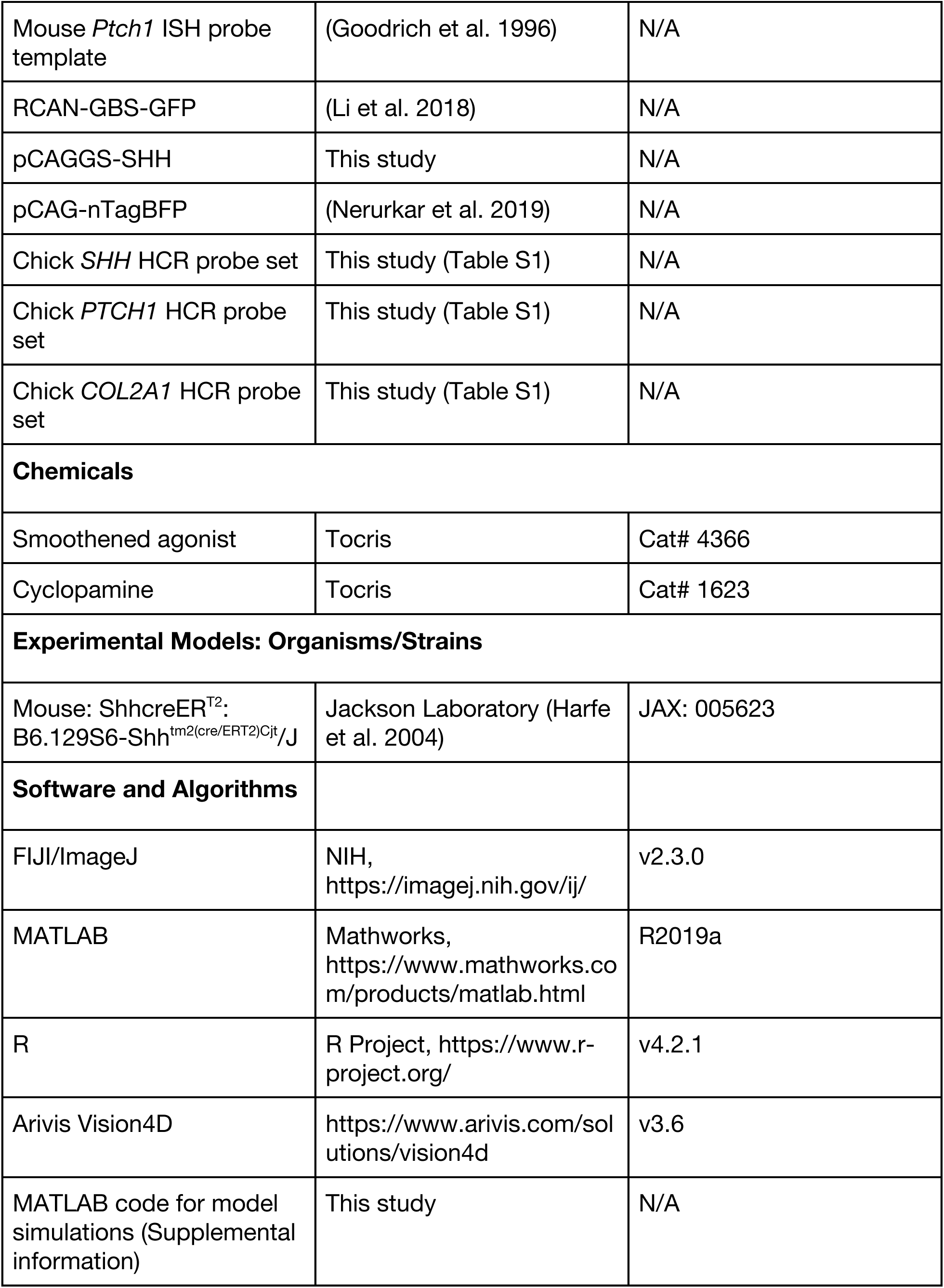

### Lead Contact and Materials Availability

Further information and requests for resources and reagents should be directed to and will be fulfilled by the Lead Contact, Clifford Tabin.

### Experimental Model and Subject Details

All animal studies were performed in compliance with NIH guidelines and standard operating protocols approved by the Institutional Animal Care and Use Committee at Harvard Medical School.

### Chicken

Fertilized Specific Pathogen Free (SPF) White Leghorn chicken eggs (Charles River Laboratories) were incubated in a 38°C chamber at 45% humidity. We staged embryos according to Hamburger and Hamilton (Hamburger and Hamilton 1951).

### Mouse

The mouse line used in this study, ShhcreER^T2^, which we refer to as *Shh-cre* in the text, has been previously described (Harfe et al. 2004). ShhcreER^T2^ heterozygous mice were provided by C. Harwell’s lab. Mice were genotyped by standard PCR (primer sequences in Table S1) using the genotyping method recommended by JAX. (https://www.jax.org/Protocol?stockNumber=005623&protocolID=30095). We crossed heterozygous mice and collected E15.5 embryos. Wild-type embryos for FISH were from timed pregnant CD-1 mice from Charles River Laboratories.

### Alligator

Alligator eggs were obtained from R. Elsey at the Rockefeller Wildlife Refuge and incubated at 32°C in moistened vermiculite, using egg band width as an approximate indicator of embryo age. Once removed from the egg, we staged embryos according to Ferguson (Ferguson 1985).

### Method Details Explant culture

Chick airways were dissected in PBS and cultured in DMEM containing 1% pen/strep and 2% chick embryo extract (US Biological). Airways were pinned to a 5% agarose base in 12-well dishes with 0.004-inch diameter tungsten rods (A-M Systems) through the larynx and through each lung, keeping the tissue taut but not stretched.

Explants were cultured near the air-liquid interface in a humidified 37°C incubator with 5% CO_2_ for the desired period. Media was removed and replaced every 48 hours. For drug treatment experiments, we diluted stock solutions of SAG (50mM in DMSO) or cyclopamine (2.5mM in ethanol) in culture media, using the equivalent volume of diluent for control explants.

### Alcian blue staining

Dissected or cultured airways were fixed overnight in 95% ethanol then transferred to staining solution (70% ethanol, 5% v/v glacial acetic acid, and 0.02% w/v Alcian blue 8GX) for 2 hours. After a brief rinse in tap water, airways were destained in 0.25% w/v KOH for 30 minutes. This was followed by clearing and destaining the stained airways 0.25% w/v KOH solutions with increasing amounts of glycerol – 25%, 33%, and 50% – each for an hour on a rocking platform. The last 0.25% KOH/50% glycerol solution was replaced and incubated on a rocking platform for at least 48 hours before imaging.

### *In situ* hybridization

For colorimetric *in situ* hybridization (ISH), fluorescent ISH (FISH), and hybridization chain reaction (HCR) labeling, dissected tissues were fixed in 4% formaldehyde/PBS for 1 hour at room temperature, then washed twice in PBS. For ISH and FISH, we transcribed DIG-labeled antisense RNA probes from previously cloned cDNA fragments (see Key Resources Table) using T3 or T7 RNA polymerases.

For ISH on sections, probes were first hybridized in the whole tissue and subsequently sectioned. Tissues were permeabilized with 10µg/ml proteinase K (NEB) for 10 minutes, refixed in 4% formaldehyde/0.2% glutaraldehyde, then hybridized overnight at 70°C with gentle shaking. Unhybridized probe was then removed with 50% formamide/2XSSC washes. Tissues were then incubated for 1 hour with blocking reagent (Sigma-Aldrich/Roche) in a maleic acid buffer and incubated with anti-DIG-AP Fab fragments (1:1000) (Sigma-Aldrich/Roche) overnight. Unbound antibody was washed out with TBS + 0.1% Tween-20, and hybridized probe was visualized with BM-purple (Sigma-Aldrich/Roche). Tissues were embedded in 10% sucrose/0.5% gelatin and sectioned at 50µm on a vibratome, then mounted for microscopy.

For FISH and HCR on sections, fixed tissues were graded through 10% sucrose and 30% sucrose into OCT. Once mounted and frozen, tissues were cryosectioned at 14µm and stored at -80°C. For FISH, DIG-labeled antisense probes were hybridized to tissue sections as described above (after (Huycke et al. 2019)), except that hybridization was done at 65°C, anti-DIG-POD Fab fragments (1:300) (Sigma-Aldrich/Roche) were bound, and probe was visualized with Tyr-Cy3 in amplification diluent (1:50) (PerkinElmer), then mounted for microscopy.

HCR probe sets were designed with R. Null’s probe generator script (https://github.com/rwnull/insitu_probe_generator) (Table S1) and ordered as oPools at 50 pmol scale (IDT). HCR on cryosections was conducted as described (Choi et al. 2018), with modifications: Tissue cryosections were washed 2 X 5 minutes with PBS and permeabilized in 70% ethanol for 1 hour, then washed 2 X 5 minutes again with PBS. Sections were then incubated in pre-warmed hybridization buffer (Molecular Instruments) for 10 minutes at 37°C. Probe pools were hybridized under plastic coverslips overnight at 37°C at 6nM in hybridization buffer. Excess probes were then removed by washing 2 X 30 minutes with probe wash buffer (Molecular Instruments) at 37°C, followed by 2 X 20-minute washes with 5X SSC + 0.1% Tween-20 (SSCT).

Sections were incubated with amplification buffer (Molecular Instruments) for 30 minutes and then incubated overnight with a solution of annealed amplifier hairpins, each at 48nM in amplification buffer, under plastic coverslips. Unbound hairpin was washed off 2 X 20 minutes with SSCT and mounted for microscopy.

### Immunostaining

For whole-mount cartilage immunostains, airways were dissected, pinned, and fixed or fixed pinned in their culture dishes with 4% formaldehyde for 1 hour at room temperature, then washed 2 X 1 hour with wash solution (PBS with 0.3% Triton X-100, 0.02% w/v SDS, and 0.2% w/v BSA). Antibodies against SOX9 (1:500, Millipore) or COL2 (1:100, DSHB) were bound overnight and unbound antibodies removed with wash solution 3 X 1 hour. Samples were then incubated overnight in species-specific fluorophore-conjugated secondary antibodies (1:500, Jackson ImmunoResearch) and subsequently washed 3 X 1 hour in wash solution before a brief rinse in PBS. Airways were then cleared for 48 hours in Sca*l*eA2 (Hama et al. 2015) before imaging.

For section immunostains, tissues were fixed in 4% formaldehyde overnight at 4°C, washed twice with PBS, then graded through 10% sucrose and 30% sucrose into OCT. Once mounted and frozen, tissues were cryosectioned at 18µm. TTF-1 staining required antigen retrieval: slides were submerged in 1X Dako Target Retrieval Solution, Citrate pH 6 (Agilent) in a glass vessel and microwaved until boiling. Once cooled, sections were then stained similarly to the whole-mount procedure described above, except that the wash solution contained 0.1% Triton X-100 and samples were not cleared. Antibodies to TTF-1/NKX2.1 and SMA were used at 1:500.

### Imaging

Micrographs of alcian blue stains were captured with a Leica M205 stereoscope with base-transmitted light. ISH sections were imaged with DIC on a Nikon Eclipse E800. Fluorescent micrographs were captured on an inverted Zeiss LSM 710 (whole-mount immunostains: Plan-Neofluar 10x/0.30 objective) or an upright Zeiss LSM 700 confocal (FISH, HCR: Plan-Apochromat 40x/1.3 Oil objective), with the pinhole set to image at 1 Airy unit. Mouse immunostains were imaged on a Nikon Ti2 inverted microscope with a W1 Yokogawa Spinning disk (50 µm pinhole), using a Plan Apo λ 20x/0.8 DIC I objective. Whole-mount immunostains were imaged in Sca*l*eA2 in a chamber constructed with clay feet under a square glass coverslip.

### Airway electroporation

For the GBS reporter experiment, eggs were incubated to HH12 (∼44 hours) and windowed to provide access to the embryo. To electroporate into the airway primordium, 2.5µg/ml solution of RCAN-GBS-GFP plasmid mixed 1:1 with 2.5µg/ml pCAG-nTagBFP electroporation control and 0.1% fast green was injected into the embryonic coelom on both sides of the embryo with a pulled glass micropipette.

Platinum electrodes were placed adjacent to the ventral (positive) and dorsal (negative) surfaces of the embryo and pulsed with five 50ms pulses of 20V supplied by a BTX electroporator. Eggs were taped and returned to the incubator until they reached HH33. Only embryos with infections throughout the airway, as assessed by 3C2 (viral gag-pro) immunostains, were included in the analysis.

For ectopic expression, the chick *SHH* CDS was cloned into a linearized pCAGGS plasmid via Gibson cloning (oligos in Table S1). A 1:1 mixture of purified pCAGGS-SHH plasmid (2.5µg/ml) and pCAG-nTagBFP (2.5µg/ml), with 0.1% fast green, was injected into the tracheal lumen of pinned airway explants with a pulled glass micropipette. Voltage was applied with a pair of platinum electrodes placed alongside the airway, using the same pulse pattern as for the coelom. For controls, mock electroporations were used. Airway explants were then cultured as described above.

### A mathematical model of tracheal cartilage patterning

Based on our experimental data, we hypothesized that tracheal cartilage patterning involves two key processes: (1) a self-organizing, Turing-like system, which generates regularly spaced stripes of cartilage; and (2) a dynamic Hedgehog gradient, which organizes the otherwise disordered Turing-like system into circumferential cartilage rings. In the following sections, we describe how we formalise these assumptions into a mathematical model, and outline the results presented in the main text.

### Model component 1: a self-organizing, Turing-like system

There are many different mechanisms that have been proposed to explain the self-organization of periodic patterns in development. The most prominent are reaction-diffusion models (Turing 1952), but mechanisms that involve other processes (such as cell movement, chemotaxis, and/or mechanical forces) can generate similar patterns *in silico* (Hiscock and Megason 2015a). Despite their differences, a common feature across all these Turing-like mechanisms is the “local activation, long-range inhibition” logic (Gierer and Meinhardt 1972; Meinhardt and Gierer 1974).

In the trachea, whilst there is good evidence that cartilage patterning is Turing-like, the precise molecular/cellular/mechanical processes that constitute this Turing system remain poorly uncharacterized (see Main Text for further discussion). Therefore, instead of assuming a specific reaction-diffusion / mechanochemical mechanism, we used a simpler and potentially more generalisable model: the Swift-Hohenberg equation (Cross and Hohenberg 1993). This equation captures the core “local activation, long-range inhibition” logic at the phenomenological level, and represents, to our knowledge, the simplest way to model a Turing instability. Moreover, previous work has shown that many of the predictions made using the Swift-Hohenberg model – including the effect of morphogen gradients – also hold true for a diverse range of Turing-like systems (Hiscock and Megason 2015a,b).

We use the Swift-Hohenberg equation to describe the time evolution of the cartilage pattern density, 𝜙(𝑥, 𝑦), via the following partial differential equation (PDE):

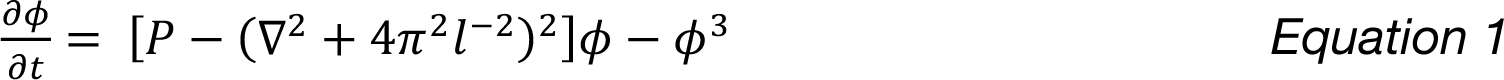

Here: 𝑃 governs the growth of the fastest-growing mode (𝑃 > 0 is required for Turing patterns to self-organize); 𝑙 denotes the spacing of the fastest growing mode; and the cubic term stabilizes the dynamics (the absence of a quadratic term favours stripes over spots).

### Model component 2: a dynamic gradient of Hedgehog signalling

#### Incorporating gradients into the model

We modified Equation 1 to incorporate the effect of a dorsal-ventral gradient in Hedgehog signalling. In general, there are two (non-mutually-exclusive) ways that a gradient could impact the Turing dynamics, resulting in the following modified Swift-Hohenberg equation:

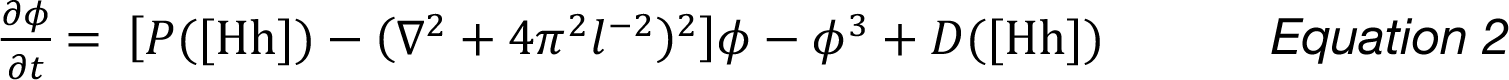

Here, [Hh] refers to the level of Hedgehog signalling activity, which modulates two terms in the equation: 𝑃([Hh]) and 𝐷([Hh]). 𝑃([Hh]) describes how Hedgehog affects the parameters of the Turing-like patterning system; hence we refer to this as a *patterning gradient* (c.f. parameter gradient in (Hiscock and Megason 2015b)). 𝐷([Hh]) describes how Hedgehog affects the production/differentiation/density of cartilage directly, independent of its effect on the Turing system; hence we refer to this as a *differentiation gradient* (c.f. production gradient in (Hiscock and Megason 2015b)).

There are a variety of molecular mechanisms that can underpin each of these terms, which also depend on the underlying Turing system. In general, we would expect that the Hedgehog gradient would not lead to a patterning or differentiation gradient alone, but rather a combination of the two together. This is because Hedgehog could interact in multiple ways with the Turing system (i.e., affecting different genes independently), and/or because the way it interacts is predicted to affect both patterning and differentiation terms simultaneously.

To capture the dorsal-ventral character of the Hedgehog gradient, we rewrite Equation 2 more explicitly as:

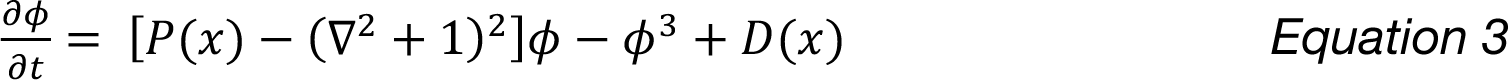

Here, 𝑥 is the circumferential spatial coordinate corresponding to the dorsal-ventral axis (see Fig 4A). Also, for simplicity we set the intrinsic cartilage spacing as a constant value throughout the simulations (𝑙 = 2𝜋)

#### Specifying the form of the gradients

The biophysical nature of Hedgehog gradient formation remains incompletely understood. Therefore, we decided to use functional forms for the gradient terms (𝑃(𝑥) and 𝐷(𝑥)), which were as simple as possible but still captured the gradient-like pattern. Based on the measured shape of the Hedgehog activity gradients and seeking a simple approximation, we therefore assumed piecewise linear functions.

We consider that the spatial variable 𝑦 varies within the range [0, 𝐿_*x*_], where 𝐿_*x*_ denotes the trachea circumference; with the ventral-most point corresponding to 𝑥 = 0, and the dorsal-most point to 𝑥 = 𝐿_*x*_/2. Because the trachea has a cylindrical geometry, this enforces periodic boundary conditions (i.e., 𝑥 = 0 is equivalent to 𝑥 = 𝐿_*x*_). Within this coordinate system, for 0 ≤ 𝑥 < 𝐿_*x*_/2, we have for the patterning gradient:

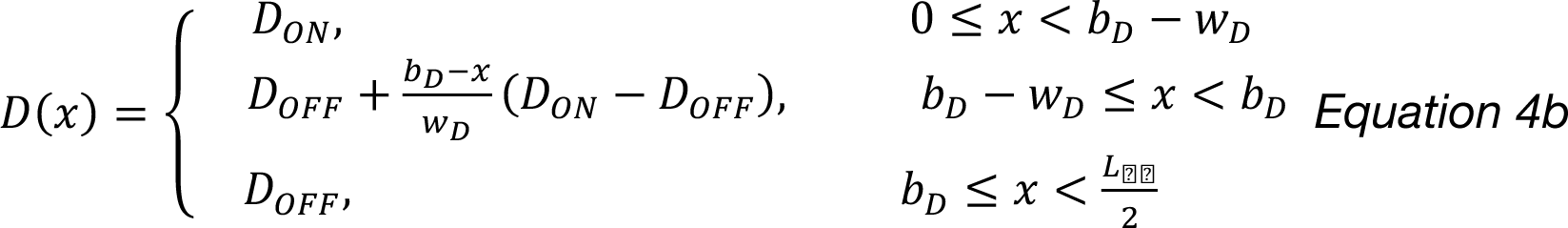

For 𝐿_*x*_/2 ≤ 𝑥 < 𝐿_*x*_, we assume that the gradient is simply the reflection of the above about the point 𝑥 = 𝐿_*x*_/2. 𝑃_*OFF*_ denotes the (saturating) minimum value of *P* [high levels of Hedgehog]; 𝑃_*ON*_ denotes the (saturating) maximum value of *P* [low levels of Hedgehog]. *P* varies linearly between these two extremes, with the linear gradient beginning at a location defined by 𝑏_*P*_ (“gradient range”), with the overall extent/width of the gradient being defined by 𝑤_*P*_ (“gradient width”). For given values of 𝑏_*P*_, 𝑤_*P*_, not all three regimes in Equation 4a may be represented (e.g., if 𝑏_*P*_ < 0 then 𝑃(𝑥) = 𝑃_(++_ everywhere).

Similarly, for the differentiation gradient:

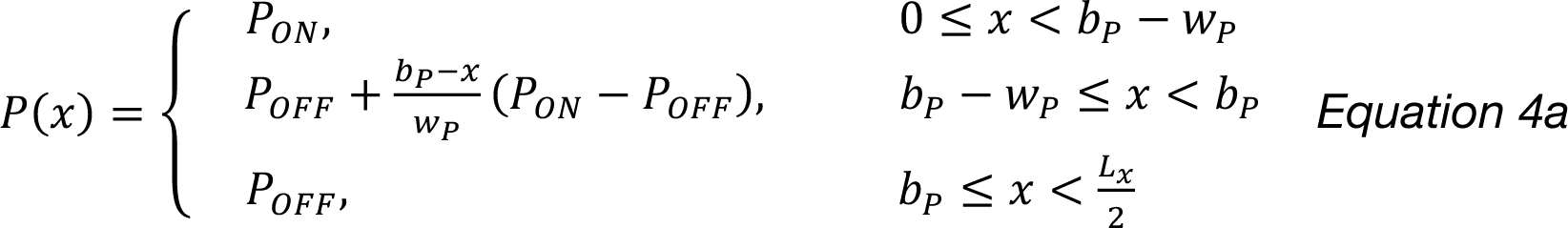

Again, for 𝐿_*x*_/2 ≤ 𝑥 < 𝐿_*x*_, we assume that the gradient is simply the reflection of the above about the point 𝑥 = 𝐿_*x*_/2.

#### Static vs. dynamic gradients

To explore the effect of these gradients on the Turing-like system, we simulated Equation 3 with these piecewise linear functions, first assuming that they were static in time. Across all parameters that we explored, we could not find any combinations of differentiation and/or patterning gradients that could reliably organize the Turing system to form circumferential cartilage rings.

Therefore, inspired by our observations of the dynamic nature of Hedgehog activity, we next investigated whether such temporally changing gradients could better organize the Turing pattern. We allowed the patterning and differentiation terms to vary in both space and time:

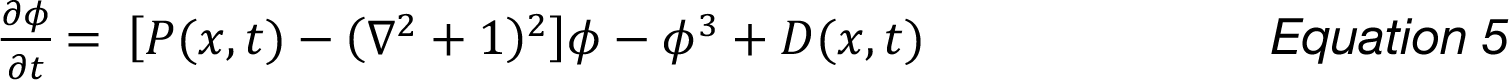

We included the temporal dynamics of Hedgehog signalling by allowing the parameters 𝑏_*P*_ and 𝑏_*D*_ to vary as a function of time:

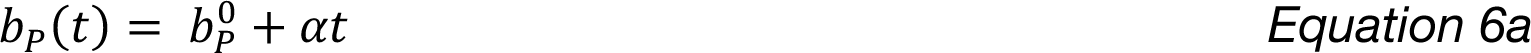

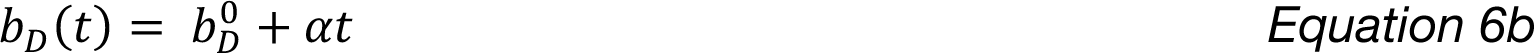

Strikingly, when we incorporated these dynamic gradients, we saw substantial alterations to the *in silico* cartilage patterns. Specifically, we found that:

*<u>Differentiation gradients alone</u>* resulted in longitudinal cartilage stripes that formed along the length of the trachea, forming sequentially in time as the gradient progressively receded towards the dorsal side.

*<u>Differentiation + patterning gradients</u>* could organize the Turing-like pattern into well-oriented cartilage stripes than spanned the entire circumference of the trachea, forming complete rings. The cartilage pattern initiates at the ventral side as a series of short, regularly spaced stripes, which then extend dorsally over time as the gradients recede.

*<u>Patterning gradients alone</u>* also generate organized cartilage rings, with similar dynamics observed with and without the differentiation gradient. Therefore, whilst a combination of differentiation and pattern gradients is likely to be operating *in vivo,* it is the presence of a patterning gradient alone which is essential to organize the pattern *in silico*.

### Generality of simulation results to other Turing-like systems

#### Theoretical arguments

Our simulations of the Swift-Hohenberg equation can explain how a dynamic signalling gradient can organize a Turing pattern into well-oriented cartilage rings. The advantage of using the Swift-Hohenberg equation is that previous theoretical work has shown that the predictions from this model also apply to a range of other Turing-like models, including reaction-diffusion systems as well as patterning based on cell movement and/or mechanical forces (Hiscock and Megason 2015b).

Based on these previous theoretical results, we predict that, regardless of parameters:

1. differentiation gradients (referred to as “production gradients” in (Hiscock and Megason 2015b)) tend to orient stripes perpendicular to the direction of the gradient
2. patterning gradients (referred to as “parameter gradients” in (Hiscock and Megason 2015b)) tend to orient stripes parallel to the direction of the gradient.
3. orientation by patterning gradients requires the gradient to transition the Turing system from below threshold (i.e., non-pattern-forming, *P < 0*) to above threshold (i.e., pattern forming *P > 0*).

Together, these theoretical properties can explain our simulation results. (1) suggests that a dorsal-ventral *differentiation gradient* would generate stripes with the wrong orientation (Fig 5C). (2) suggests that a dorsal-ventral *patterning gradient* would be required to produce stripes with the correct circumferential orientation. (3) suggests that significant orientation effects occur only when the *patterning gradient* defines a transition/boundary between patterning-forming and non-pattern-forming regions.

Together, (2) and (3) suggest that a static patterning gradient cannot form well-oriented rings, because in this case the whole circumference would be pattern forming and therefore in the absence of a transition/boundary region, orientation effects would be minimal (Fig 5B). However, by having a dynamic patterning gradient, this can allow well-oriented but short stripes to initiate at the ventral regions (Fig 4C). Over time, the gradient dynamics allows the pattern-forming region to expand dorsally, with stripes continuing to form along the gradient (dorsal-ventral) direction (Fig 4C). This continues until the pattern forming domain covers the entire circumference and complete rings have formed (Fig 4C).

Because the results (1-3) appear to be a general property of Turing patterns, we argue that our findings will likely generalize to other Turing-like systems, including the (as yet unidentified) mechanism that governs tracheal cartilage patterning.

#### A reaction-diffusion model with dynamic gradients

In addition to these theoretical arguments, we investigated a reaction-diffusion model to explore whether our findings from the Swift-Hohenberg equation can generalize to other Turing-like models. We assume that tracheal cartilage patterning is controlled by a network of two interacting and diffusing secreted molecules, which we term activator and inhibitor respectively (Fig S4A). The activator positively autoregulates itself, as well as inducing its own (more rapidly diffusing) inhibitor; a regulatory logic that is commonly found in reaction-diffusion mechanisms of periodic patterning (Kondo and Miura 2010). For the precise functional forms of these interactions, we take inspiration from the CIMA chemical reaction – one of the first experimentally realized Turing patterns (Lengyel and Epstein 1991, 1992). We modify the classic CIMA reaction-diffusion equations to introduce a patterning gradient within the system, specifically:

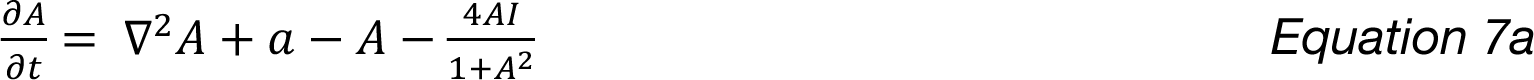

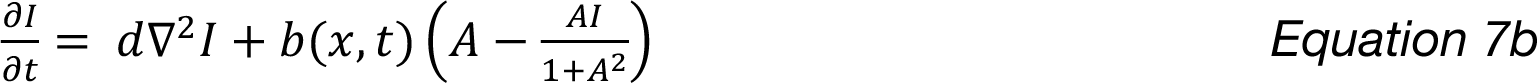

Here, 𝑏(𝑥, 𝑡) represents a dynamic, dorsal-ventral patterning gradient; we assume piecewise linear functions, defined similarly to Equations 4&5. When we simulate a receding gradient in 𝑏, this organizes the Turing stripes into well-oriented circumferential rings, with similar dynamics to the Swift-Hohenberg model (Fig S4C).

Whilst we do not speculate here on putative activator(s) or inhibitor(s) involved in cartilage patterning, these results provide a proof-of-principle demonstrating that dynamic gradients can organize reaction-diffusion Turing patterns similar to the predictions made by the Swift-Hohenberg model.

### Simulation methods

Each of our models represents a dynamical system defined by a PDE, or system of PDEs (static gradients: Equation 3; dynamic gradients: Equation 5; CIMA reaction: Equation 7). We solve these PDEs using custom MATLAB scripts. All scripts and parameter values used in this study are provided as supplemental materials.

Briefly, we assume a cylindrical geometry as schematized in Fig 4A, with the 𝑥-axis referring to the azimuthal/circumferential coordinate (ranging from *0* to 𝐿_*x*_) and the *y*-axis referring to the longitudinal axis (ranging from *0* to 𝐿_6_). We employ a pseudo-spectral implicit-explicit numerical scheme to solve the stiff PDEs. Briefly, we discretize the space into a 𝑁_*x*_ × 𝑁_6_ grid, which transforms the PDEs into a system of ODEs. At each time step, we combine the (implicit) backward Euler algorithm with the Fast Fourier Transform (FFT) algorithm to efficiently compute the increments associated with the spatial derivative terms in the PDEs [(∇^$^ + 1)^$^𝜙 for Equations 3&5; ∇^$^𝐴 and ∇^$^𝐼 for Equation 7). We then use the (explicit) forward Euler algorithm to compute the increments associated with all remaining terms. The gradient terms 𝑃(𝑦, 𝑡), 𝐷(𝑦, 𝑡), 𝑏(𝑦, 𝑡) are represented by matrices, whose values are determined by Equations 4&6. Initial conditions are set by adding normally distributed random numbers to a uniform state.

We approximate the geometry of the trachea as cylindrical. Given that a cylinder has zero Gaussian curvature, this means that the Laplacian terms can be computed assuming a flat, 2D geometry (Faraudo 2002; Yoshigaki 2007). However, in contrast to a 2D sheet, we must assume periodic boundary conditions for the *x*-axis; this is automatically encoded in the FFT algorithm that we use. For the *y*-axis boundary conditions (at the anterior/posterior boundary of the trachea), we consider two cases:

1. *Reflective boundary conditions:* We assume reflective boundary conditions at both anterior and posterior boundaries.
2. *“Repressive” boundary conditions:* We assume that 𝑃 = 𝑃_78#9:;<_ < 0, 𝐷 = 𝐷_78#9:;<_ < 0 for all regions above the anterior boundary and below the posterior boundary.

To incorporate these boundary conditions, we extend the simulation domain a length, 𝑙_*b*_, at both anterior and posterior ends, and refer to these regions as the *boundary regions*. We follow the approach taken in (Macdonald et al. 2013): for reflective boundary conditions, we project the value of 𝜙 outwards from the boundary to fill the entire boundary region; for repressive boundary conditions, we set 𝑃, 𝐷 as defined above in the boundary regions which prevents patterning.

We found that *reflective boundary conditions* led to mild defects in pattern orientation near the anterior/posterior boundaries (Fig S2B). Therefore, we used *repressive boundary conditions* for all other figures, which displayed more robust pattern orientation. When inspecting the dynamics of *in silico* ring formation, we saw an hour-glass-like effect, in which cartilage stripes initiated slightly earlier towards the anterior and posterior boundaries, and for a short period were initially longer in these regions (Fig 4D). We saw no such hour-glass pattern when using *reflective boundary conditions* (Fig S2A). Therefore, we propose that some form of *repressive boundary conditions* must be operating *in vivo,* although we do not speculate as to their molecular implementation.

We found that local pre-patterns of cartilage that mimic the avian syrinx could generate similar hour-glass-like patterns (Fig 4D; here we modify 𝐷 to promote 2 large bands of cartilage). These patterns were absent in the corresponding region of the syrinx-less alligator trachea (Fig S2B). These results further support the idea that localized changes along the anteroposterior axis can lead to boundary effects resulting in transient hour-glass-like patterns.

#### Modelling the effect of Hedgehog perturbations

To mimic the effect of persistent Hedgehog activity following electroporation, we introduced a term 𝑃_<’7=<>789_(𝑥, 𝑦). For simplicity, we assume that this function is radially symmetry about the electroporation point (𝑥_0_, 𝑦_0_), i.e., can be written as a function 𝑃_<’7=<>789_(𝑟) where 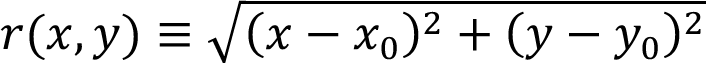. We assume a similar pattern of activity as in Equation 4a:

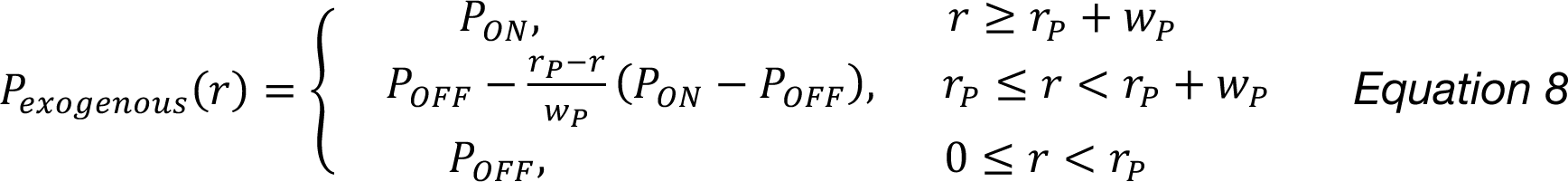

Here, 𝑟_*P*_ plays an analogous role to 𝑏_*P*_ as in Equation 7a, but with the crucial distinction that 𝑟_*P*_ is constant in time. To combine the exogenous, constant source of Hedgehog (Equation 8) with the endogenous, dynamic source of Hedgehog (Equation 4), we assume that their combined action is defined by the elementwise minimum of the two terms. We define 𝐷_<’7=<>789_ in the same manner; we set 𝑟_*D*_ − 𝑟_*P*_ = 𝑏^0^_*D*_ − 𝑏^0^_*P*_ for consistency.

We also modelled the addition of cyclopamine to the *ex vivo* tracheas, applied at time 𝑇_*c*_. Before the perturbation, i.e., 𝑡 ≤ 𝑇_*c*_, we set 𝑃, 𝐷 via Equation 4. After the perturbation, i.e., 𝑡 > 𝑇_*c*_, we set 𝑃 = 𝑃_*ON*_, 𝐷 = 𝐷_*ON*_, assuming that the inhibition of Hedgehog activity is fast relative to the patterning kinetics.

#### Exploring pattern defects

In contrast to the straight circumferential rings in the chick, cartilage patterns in mouse/human trachea are more disordered with various types of defects present. We wondered whether the patterns of these defects could be predicted by small modifications to our chick-inspired model. For the sake of simplicity, we focused on the scenario where Hedgehog generates a patterning gradient only, because it is this term which is essential to organize the pattern into circumferential rings (Fig 5C).

First, we varied the width of the gradient 𝑤_*P*_ (Fig 5A). We found that cartilage rings reliably formed across a large range of gradient widths. This included, at one extreme, gradients with a width comparable to the trachea circumference; to the other extreme, in which the gradients were narrow and could be approximated as step-functions. This demonstrates that, as long as there is a boundary between pattern-forming and non-pattern-forming regions, the resulting stripes will form perpendicular to this boundary, mirroring the theoretical arguments above. In the limit where the gradient width is very narrow, the gradient dynamics lead to a “wave of competency” which sweeps from ventral to dorsal across the trachea; similar waves of competency have been proposed to be involved in patterning the avian skin (Bailleul et al. 2019; Ho et al. 2019), and lead to some non-intuitive effects on reaction-diffusion models (Liu et al. 2022).

Second, we varied the gradient parameters 𝑃_*ON*_, 𝑃_*OFF*_. For simplicity, we set 𝑊 = 0 for these comparisons. We found that cartilage rings reliably formed for a wide range of 𝑃_()_values. In contrast, we found that increasing the value of 𝑃_(++_ led to more disorganized patterns (Fig. 5A). For intermediate values of 𝑃_(++_, we observed the formation of cartilage rings, but with a variety of ring defects present. Similarly, we varied the parameter 𝑏^0^, which can be interpreted as the initial *range* of Hedgehog activity. For low values of 𝑏^0^ (i.e., Hedgehog signalling is initially high across the entire dorsoventral axis), straight cartilage rings formed. Whereas for higher values of 𝑏^0^ (i.e., Hedgehog signalling begins lower in the ventral regions), this leads to pattern defects and disorganized stripes (Fig 5A).

Finally, for an intermediate value of 𝑏^0^, we repeated the simulations with the same parameters but with different (noisy) initial conditions. We saw a variety of defect types that are reminiscent of those observed in human and mouse trachea (Fig S3). These included: branched rings; incomplete rings; regional thickenings; and H-, X- or Z-shaped elements, all of which have been reported in mammalian trachea (Vanpeperstraete 1973; Sala et al. 2011; Premakumar et al. 2018; Lam et al. 2020).

Whilst we do not consider these simulations to be an accurate model of mouse/human tracheal patterning (because, for example, they neglect the essential but poorly understood role of smooth muscle), they do show that an intrinsically Turing-like mechanism, when coupled to a dynamic signalling gradient, can generate pattern defects that resemble the variations of cartilage patterns in these other species, further lending support to our model.

## SUPPLEMENTAL INFORMATION

**• Figures S1–S5**

Fig. S1. Tracheal geometry and growth rates are not sufficient to orient the tracheal ring pattern.

Fig. S2. Boundary conditions can produce edge effects resembling early chondrogenesis.

Fig. S3. Varying initial simulation conditions produces irregularities resembling those occurring in tracheal cartilage.

Fig. S4. A reaction-diffusion model produces similar results to the general patterning model.

Fig. S5. *Shh* is required for dorsal smooth muscle formation in mouse.

- **Table S1. Oligonucleotide sequences**
- **MATLAB code for model simulations**

## ACKNOWLEDGEMENTS

We are grateful to the members of the Tabin and Cepko labs for helpful discussions throughout this study and especially to ChangHee Lee for his help with mouse breeding during pandemic lockdown. Corey Harwell and Steve Vu provided ShhcreER^T2^ mice, and Ruth Elsey and the Rockefeller Wildlife refuge provided fertilized alligator eggs. We also thank the Microscopy Resources on the North Quad (MicRoN) core at Harvard Medical School. And we are grateful to Michael Dyer, his lab, and St. Jude DNB for generously hosting EPK during much of this project. This work was funded by a grant from the Gordon and Betty Moore Foundation Grant 4498 to CJT.

## Author Contributions

EPK and CJT conceived the study. DGM crossed, staged, genotyped, immunostained, and imaged Shh-cre mice. EPK conducted the other experiments. TWH designed and conducted modeling and simulations. EPK, CJT, and TWH wrote the manuscript with input from DGM.

