## Supplemental figures for "A dynamic Hedgehog gradient orients tracheal cartilage rings"

### SUPPLEMENTAL FIGURE LEGENDS

**Figure S1. Tracheal geometry and growth rates are not sufficient to orient the tracheal ring pattern. A.** Simulations on cylinders of varying diameter showing that extreme geometries are required to orient circumferential rings, such that the tube circumference is on the order of, or smaller than the pattern wavelength, which is not true for the trachea. **B.** Ratios of the tracheal circumference to the anteroposterior distances between cartilage rings for chick (HH35) and mouse (E13.5) show that tracheal geometry alone is insufficient to orient the airway cartilage pattern.

**Figure S2. Boundary conditions can produce edge effects resembling early chondrogenesis. A.** Simulations with reflective boundary conditions at the anterior/posterior ends do not show a transient hour-glass pattern (left) and display some non-oriented patterns at the ends of the cylinder (right). **B.** Simulating cartilage patterning without a posterior boundary (present in Fig. 4D) produces a ventral tapered domain of initial chondrogenesis, but only at the anterior end of the trachea. Here, we assume that cartilage is also being patterned more posteriorly than is shown. This continuation of patterning beyond the posterior extent of the trachea and into the tracheobronchial junction mimics the assumed state of signaling in taxa lacking a syrinx. **C.** Alcian-blue-stained alligator airways, which lack a syrinx, show a tapered domain of initial chondrogenesis at the ventral anterior surface of the trachea, in contrast to the initial pattern in the chick airway (Fig. 4E).

**Figure S3. Varying initial simulation conditions produces irregularities resembling those occurring in tracheal cartilage.** Fifteen examples of model simulations with different initial conditions, showing a variety of pattern irregularities that resemble those present in tracheas (compare with Fig. 3) (Lam et al., 2020; Premakumar et al., 2018; Vanpeperstraete, 1973).

**Figure S4. A reaction-diffusion model produces similar results to the general patterning model. A.** Top: schematic of the two-species reaction-diffusion system. Bottom: PDE model describing the reaction-diffusion model (see Methods) with a term for the Hh gradient effect. **B.** Simulation of a reaction-diffusion model without a gradient effect produces a disorganized pattern. **C.** Simulation of a reaction diffusion model with a retracting gradient produces a similar pattern evolution and final oriented pattern as the simpler Swift-Hohenberg model (Fig. 4C).

**Figure S5. *Shh* is required for dorsal smooth muscle formation in mouse.** Immunostains (SMA = smooth muscle marker; NKX2.1 = respiratory epithelium marker) on transverse sections of E15.5 littermates. **A.** Heterozygous-wild-type mice have normal smooth muscle (white arrowhead) at the dorsal aspect of the trachea (3 of 3 embryos). **B.** Homozygous mutant mice lack respiratory smooth muscle (empty arrowhead) (3 of 3 embryos). Scale bars = 100μm; es = esophagus; tr = trachea.

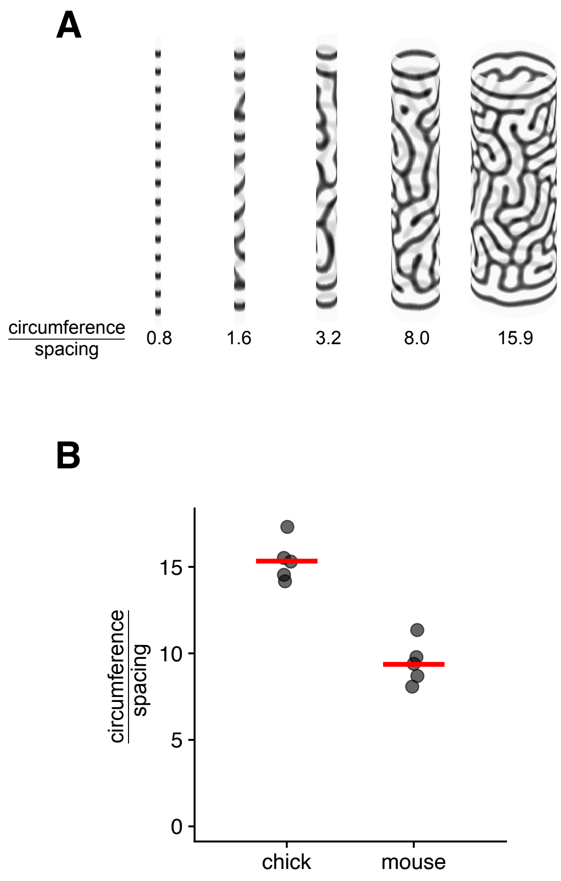
A
circumference / spacing
0.8
1.6
3.2
8.0
15.9
B
circumference / spacing
0
5
10
15
chick
mouse

Figure S1

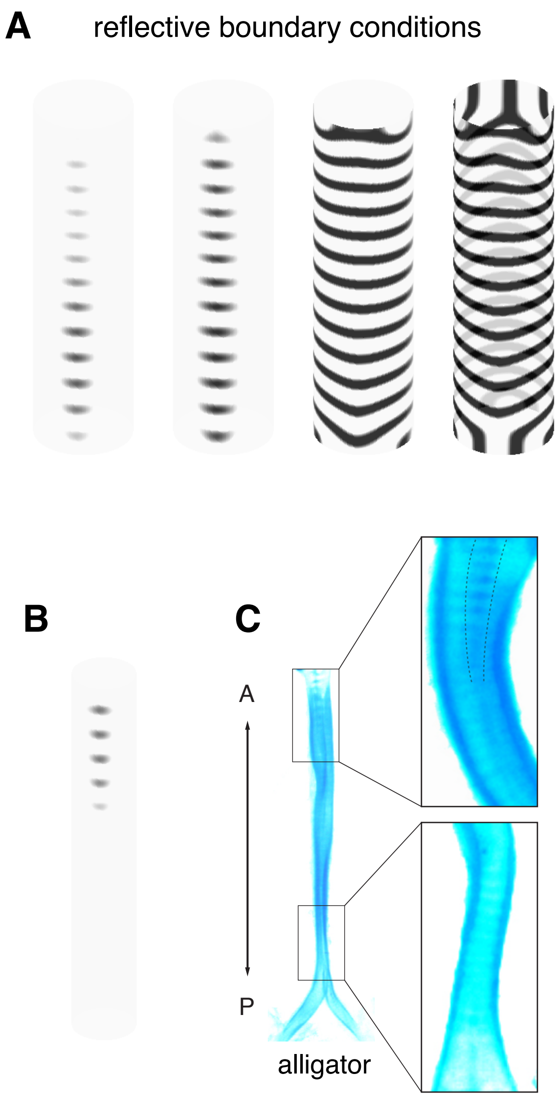


Figure S2

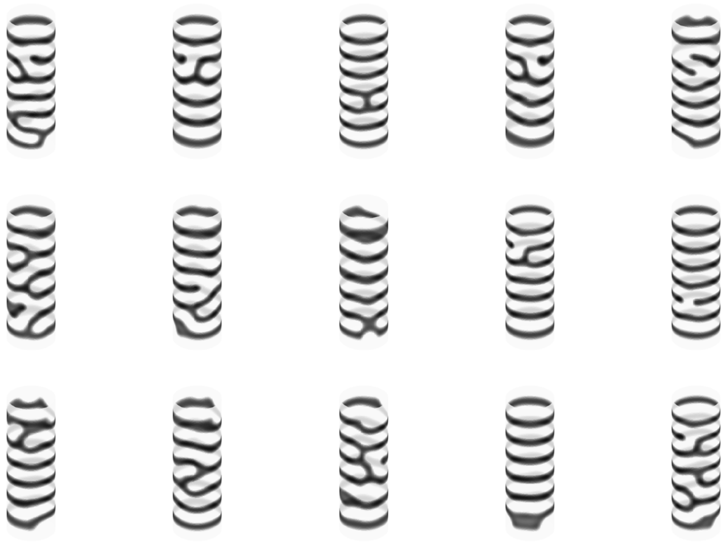

**simulation results with varied initial conditions**

Figure S3

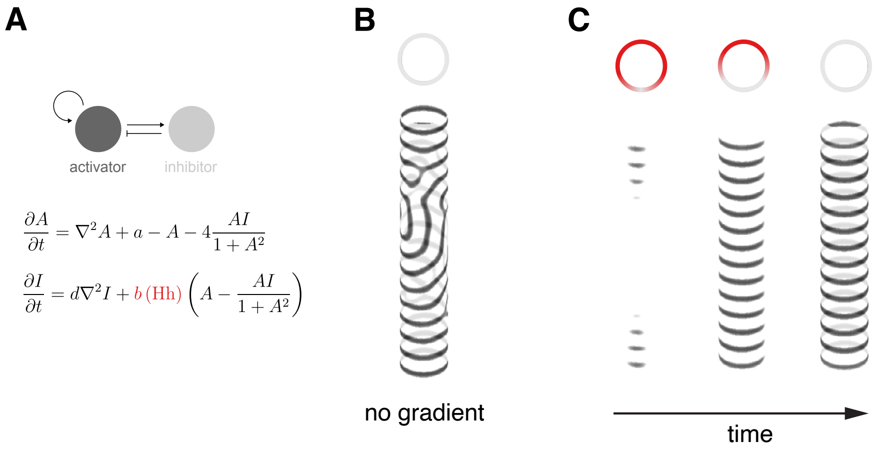
A
B
C
activator
inhibitor
$\frac{\partial A}{\partial t} = \nabla^2 A + a - A - 4\frac{AI}{1+A^2}$
$\frac{\partial I}{\partial t} = d\nabla^2 I + b\,(\mathrm{Hh})\left(A - \frac{AI}{1+A^2}\right)$
no gradient
time

Figure S4

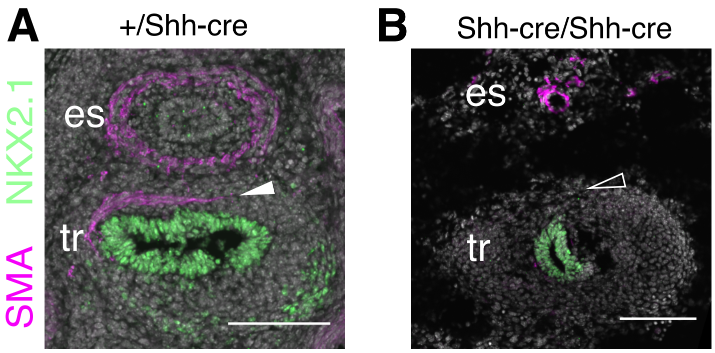


Figure S5
